# Giraffe Translocation Population Viability Analysis

**DOI:** 10.1101/619114

**Authors:** Derek E. Lee, Elmar Fienieg, Cock Van Oosterhout, Zoe Muller, Megan Strauss, Kerryn D. Carter, Ciska P.J. Scheijen, Francois Deacon

## Abstract

Most populations of giraffes have declined in recent decades, leading to the recent decision to upgrade the species to vulnerable status, and some subspecies to endangered. Translocations have been used as a conservation tool to re-introduce giraffes to previously occupied areas or establish new populations, but guidelines for founding populations are lacking. To provide general guidelines for translocation projects regarding feasibility, we simulated various scenarios of translocated giraffe populations to identify viable age and sex distributions of founding populations using Population Viability Analysis (PVA) implemented in Vortex software. We explored the parameter space for demography (population growth rates: λ = 1.001, 1.010, 1.024), and the genetic load (number of lethal equivalents: LE = 2.5, 6.29, 12.6), examining how variation in founding numbers (N = 5 to 80 females) and sex ratios (M:F = 0.1 to 0.5) affected 100-year probability of extinction and genetic diversity. We found that even very small numbers of founders (N ≤10 females) can appear to be successful in the first decades due to transient positive population growth, but with moderate population growth rate and moderate genetic load, long-term population viability (probability of extinction <0.01) was only achieved with ≥30 females and ≥3 males released. To maintain >95% genetic diversity of the source population in an isolated population, 50 females and 5 males are recommended to comprise the founding population. Sensitivity analyses revealed first-year survival and reproductive rate were the simulation parameters with the greatest proportional influence on probability of extinction and genetic diversity. These simulations highlight important considerations for translocation success, and data gaps including true genetic load in wild giraffe populations.

---

*Giraffes (Giraffa camelopardalis*) are endemic African ruminant ungulates, and one of only a handful of extant terrestrial megaherbivores (Owen-Smith 1988). Most populations of giraffes have declined in recent decades, leading to the recent decision to upgrade the species to vulnerable status, and some subspecies as endangered, on the IUCN Red List (Muller et al. 2018). Translocations have been used as conservation tools to re-introduce giraffes to previously occupied areas or establish new populations (Winter et al. 2019), but quantitative guidelines for establishing viable populations of translocated giraffes do not exist.

General guidelines for translocations suggest population modelling should be used to assist in the assessment of project feasibility (IUCN 2013). Population modelling for translocation projects can offer guidance on age- and sex-classes for founder populations by providing estimates of population persistence, long-term viability, and genetic diversity (Seddon et al. 2007). Population viability analysis (PVA) is commonly used to predict the likely future status of a population and thus offers a quantitative basis for evaluating alternative management strategies (Boyce 1992, Morris and Doak 2002). The availability of software programs such as Vortex has facilitated the application of population modelling in re-introduction planning (e.g., Bustamante 1998) and post-release evaluation (e.g., Slotta-Bachmayret et al. 2004).

Our objective here is to identify the minimum size and sex ratio of a founding population that would ensure long-term population viability and genetic diversity. We simulated various scenarios of translocated giraffe populations using PVA implemented in Vortex software (Lacy and Pollack 2017). We used stochastic, single-population, individual-based models to project future population trajectories and estimate probability of extinction (PE), stochastic rate of population increase (r), and genetic diversity (GD) at a hypothetically ideal prospective release site under different founder population release scenarios. Scenarios varied the numbers of females and males to release (Converse et al. 2013) and included 3 levels of inbreeding genetic load under demographic rates simulating slow, moderate, and fast population growth rates. We considered 4 criteria for defining successful translocations: 100-year probability of extinction PE < 0.05 and PE < 0.01 (Morris and Doak 2002); and 100-year genetic diversity GD > 80% and GD > 95% of the source population (Frankham et al. 2010). We also performed a sensitivity analysis to determine which demographic and genetic load parameters had the greatest proportional influence on PE and GD.

## METHODS

### PVA Scenarios

We constructed population models in program Vortex to simulate a hypothetical translocated population of simulated individuals using demographic parameters from literature (Lee and Strauss 2016, Dagg 2014) and publications of the Giraffe International Studbook (Lackey 2009). Males bred from ages 2 to 25, and females bred from ages 3 to 29, the maximum age of survival for both sexes was 30 years, females always produced 1 calf, and sex ratio at birth was equal (Dagg 2014). Within the model parameters, offspring were dependent upon their mother for 1 year, meaning that if the mother died during the calf’s first year, then the calf also died (Dagg 2014). All males aged 2 and above were in the breeding pool. Demographic rates are given in **Table 1**.

**Table 1.** Demographic rates and variability (SD = standard deviation) used to parameterize population simulations of giraffe translocations reflecting slow, moderate, and fast population growth rates.

| Population growth rate | <u>Slow</u> |  | <u>Moderate</u> |  | <u>Fast</u> |  |
| --- | --- | --- | --- | --- | --- | --- |
| $\lambda$ | 1.001 | | 1.010 | | 1.024 | |
| Demographic rates | % | (SD) | % | (SD) | % | (SD) |
| Mortality Age 0-1 | 50 | (10) | 50 | (10) | 40 | (10) |
| Mortality Age 1-2 | 20 | (6) | 20 | (6) | 20 | (6) |
| Mortality Age >2 | 5 | (3) | 5 | (3) | 5 | (3) |
| Reproduction | 45 | (10) | 50 | (10) | 50 | (10) |

To parameterize our PVA, we based our demographic rates on published observations of means and variances of age-specific survival and fecundity for wild giraffe populations throughout Africa (Lee and Strauss 2016), and data for reproductive longevity and inbreeding depression from the global zoo population (Lackey 2009). We used data from IUCN Red List assessments to compute mean population growth rates for 7 growing giraffe subpopulations, including some translocations (Muller et al. 2018). Mean observed population growth rate was 1.024 (range 1.0045 to 1.035). Because the highest observed growth rates were from short time spans, they were likely due to transient population dynamics and not the asymptotic growth rate. Therefore, in our PVA we simulated populations with 3 different demographic parameterizations with different asymptotic population growth rates (λ) that represented slow- (1.001), moderate- (1.010), and fast-growing (1.024) populations (Lee and Strauss 2016, Muller et al. 2018). The fastest population growth rate is probably only achievable in translocation destinations that have few or no large predators (Lee and Strauss 2016, Muller et al. 2018).

Given that few individuals typically are translocated on any given occasion, demographic stochasticity, Allee effects, and inbreeding depression could all adversely affect long term viability for translocated populations (O’Grady et al. 2006, Deredec and Courchamp 2007, Van Houtan et al. 2010). We included stochasticity (demographic rates vary among individuals and over time) and inbreeding depression (reduced reproduction and calf survival due to inbreeding) in our simulations because these effects are likely to exist in giraffe populations (O’Grady et al. 2006, Lackey 2009, Lee and Strauss 2016, Lacy and Pollack 2017). We did not include Allee effects (reduced demographic rates at small population size) because Allee effects are unlikely in giraffe populations, especially in enclosed reserves where mates can easily locate each other.

We included 3 levels of inbreeding depression. Using data from the giraffe studbook, we estimated the minimum genetic load as number of total lethal equivalents (LE) per individual with a regression analysis of natural-log-transformed first-year survival on F-coefficient according to Morton et al. (1956). The slope of the regression provided an estimate of the reduction of fitness due to inbreeding and gave the lower bound approximation of the effective number of LE per gamete, so we doubled the slope to estimate the low value for number of LE per zygote (Lynch and Walsh 1998). We used Vortex inbreeding values representing a low genetic load of 2.5 LE per individual (from our regression), a moderate load of 6.29 LE (O’Grady et al. 2006, Nietlisbach et al. 2018), and a high genetic load of 12.6 LE (double the moderate load). Fifty percent of LE are due to recessive lethal alleles. The other half of the LE are due to overdominance (heterozygote superiority), and this genetic load cannot be removed by selection (Lacy and Pollack 2017). We set demographic temporal correlation between mortality and reproduction due to environmental variation at 0.5.

We assumed zero translocation-related mortality, so no additional mortality effect above normal levels due to the process of capturing, relocating, and releasing. If mortality is expected during the translocation process, our simulation results should be interpreted using the actual number of successful live releases. We assumed zero post-release dispersal movements because many translocations will likely be into fenced or otherwise constrained areas, and because we wanted to keep track of every individual in the translocated population.

We projected 198 PVA scenarios. We simulated populations with various numbers of 2-year-old females released (5, 10, 20, 30, 40, 50, 60, and 80 females) and different numbers of 2-year-old males released to vary the sex ratio (SR) at release (SR = males / females, range = 0.1 to 0.5). We simulated each combination of number of females and males at the 3 levels of asymptotic growth rate, and 3 levels of genetic load.

For all scenarios we ran 1000 iterations of our stochastic model to project the populations for 100 years. We selected a projection time of 100 years because giraffes are a long-lived species and we were most interested in the longer-term implications of translocation decisions, particularly the effect of the number of individuals released. Extinction definition was N < 2. We assumed a hypothetically ideal release site, but included effects of density dependence with a carrying capacity of N = 1000. This was done because we were primarily interested in the effects of demography and genetics without the confounding effects of habitat limitation, which varies considerably between translocation sites. Forage quality and availability in all seasons should be quantified at the release site to determine carrying capacity as part of a comprehensive pre-translocation assessment. For each scenario we recorded output on PE, and GD (as observed heterozygosity) in the final extant population. We provide guidelines for founding population size and sex ratio for successful translocations with success defined 4 ways: probability of extinction PE < 0.05 and PE < 0.01; and genetic diversity GD > 80% and GD > 90%.

We performed a sensitivity analysis to determine which demographic or inbreeding parameter was most influential to long-term viability by comparing outcomes from simulations that were identical in every way except that we reduced a single demographic or inbreeding parameter by approximately 25%. We simulated release of 30 females and 3 males using moderate demographic rates (Table 1, middle columns) and moderate genetic load (LE = 6.29) as the reference simulation. We then simulated 6 identical populations except each had a single change: 1) increased age of first reproduction by 1 year for males and females; 2) first year mortality = 63%; 3) second year mortality = 25%; 4) adult mortality = 6%; 5) reproduction = 38%; and 6) LE = 7.9. The observed differences in PE and GD between the changed simulations and the reference simulation indicates the sensitivity due to a proportionally similar change in each demographic or genetic load parameter.

## RESULTS

Population projections from all our scenarios followed the same general post-release trajectory. Overall, there was an initial population decline for 1 year as some released individuals died before maturity, followed by 27 years of rapid population growth as the founders and their offspring reproduced, then a 1-year decline in year 29 as the last remaining founding animals died. This was followed by a brief 3-year period of stable or positive population growth until the first post-founder generation died and stable age distribution was achieved. Long-term (100-year) viability depended on population growth rate and inbreeding genetic load, as well as which success criteria were used, but minimum founding population varied from 10 to 60 females and 1 to 15 males (**Table 2**).

**Table 2.** Minimum founding population sizes of female (F) and male (M) giraffes for translocations to achieve 100-year population viability success under 4 criteria of success, with 3 levels of intrinsic population growth rate and 3 levels of inbreeding genetic load.

| Criteria | Population growth rate | Inbreeding genetic load (LE = lethal equivalents) |  |  |
| --- | --- | --- | --- | --- |
|  |  | Low (2.5 LE) | Mod (6.3 LE) | High (12.6 LE) |
| <0.05 Probability of Extinction | Slow (1.001) | 20F 2M | 30F 3M | 30F 6M |
|  | Mod (1.010) | 20F 2M | 20F 2M | 30F 3M |
|  | Fast (1.024) | 10F 3M | 10F 5M | 20F 2M |
| <0.01 Probability of Extinction | Slow (1.001) | 30F 3M | 30F 6M | 40F 5M |
|  | Mod (1.010) | 20F 2M | 30F 3M | 30F 9M |
|  | Fast (1.024) | 10F 5M | 20F 2M | 20F 5M |
| >80% Genetic Diversity | Slow (1.001) | 20F 2M | 20F 2M | 30F 3M |
|  | Mod (1.010) | 20F 2M | 20F 2M | 20F 2M |
|  | Fast (1.024) | 10F 1M | 10F 3M | 20F 2M |
| >95% Genetic Diversity | Slow (1.001) | 50F 15M | 60F 12M | 60F 18M |
|  | Mod (1.010) | 50F 5M | 50F 5M | 50F 10M |
|  | Fast (1.024) | 40F 5M | 40F 5M | 40F 5M |

The 100-year PE declined as the number of released females, and sex ratio increased (**Fig. 1**). Assuming fast population growth demographic rates and minimum inbreeding genetic load, the 100-year PE was below 0.05 when release included at least 10 females and 3 males (**Table 2**). Under more conservative demographic rates (λ = 1.001) and realistic inbreeding genetic loads (LE = 6.3 or 12.6), the minimum release population for 95% viability at 100 years was 30 females and 3–6 males (**Table 2**). Final GD increased with larger numbers of females and males in the founding population (**Fig. 2**). To meet the success criteria of preserving 80% of GD, minimum founding population sizes were similar to those required for the criteria of PE < 0.05, i.e. 30 females and 3–6 males (**Table 2**).

**Figure 1.**
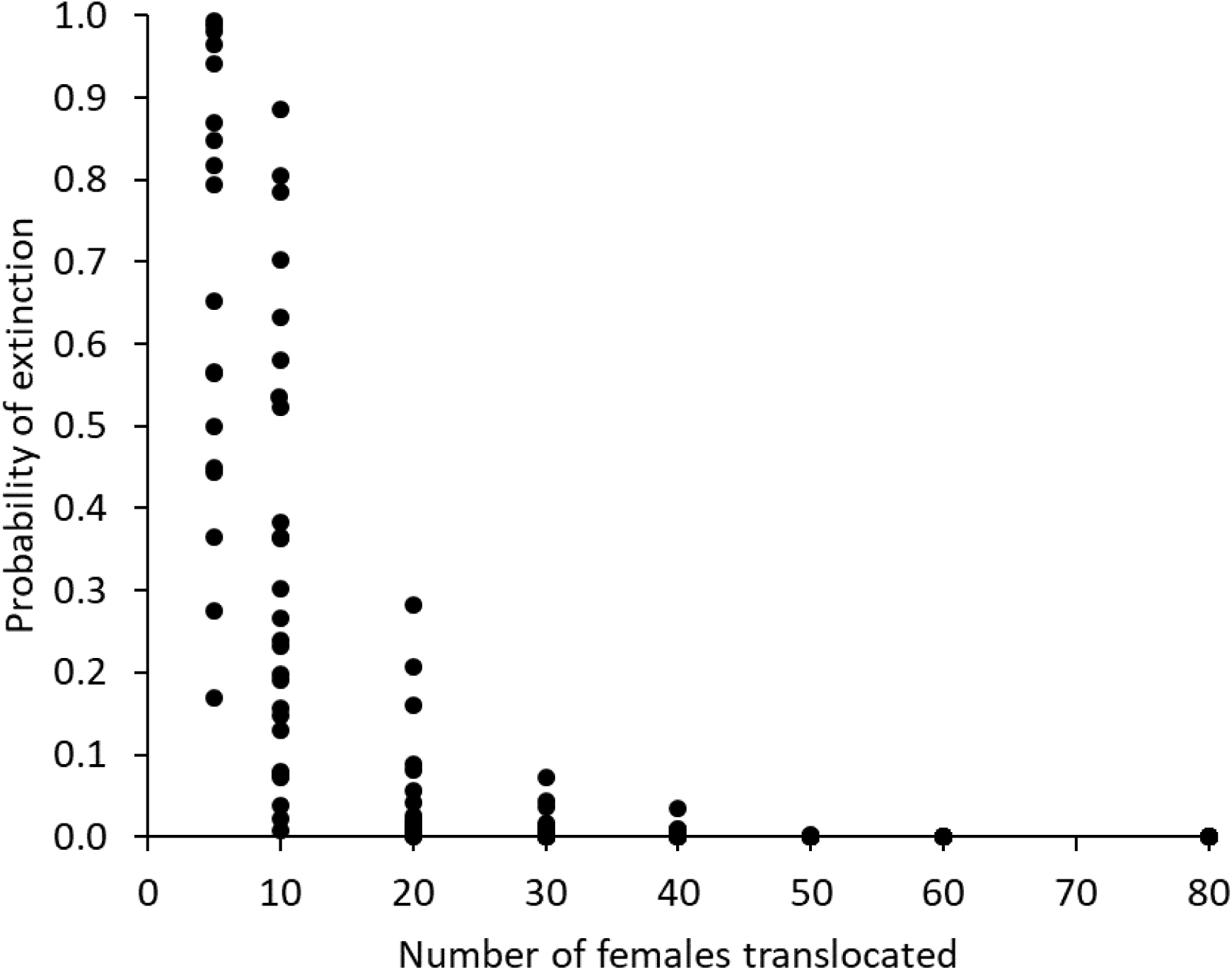
100-year probability of extinction decreased as the number of released females increased in population projections simulating translocations of different numbers of wild giraffes. Simulations used population growth rates: λ = 1.001, 1.010, 1.024; and genetic load number of lethal equivalents: LE = 2.5, 6.29, 12.6.

**Figure 2.**
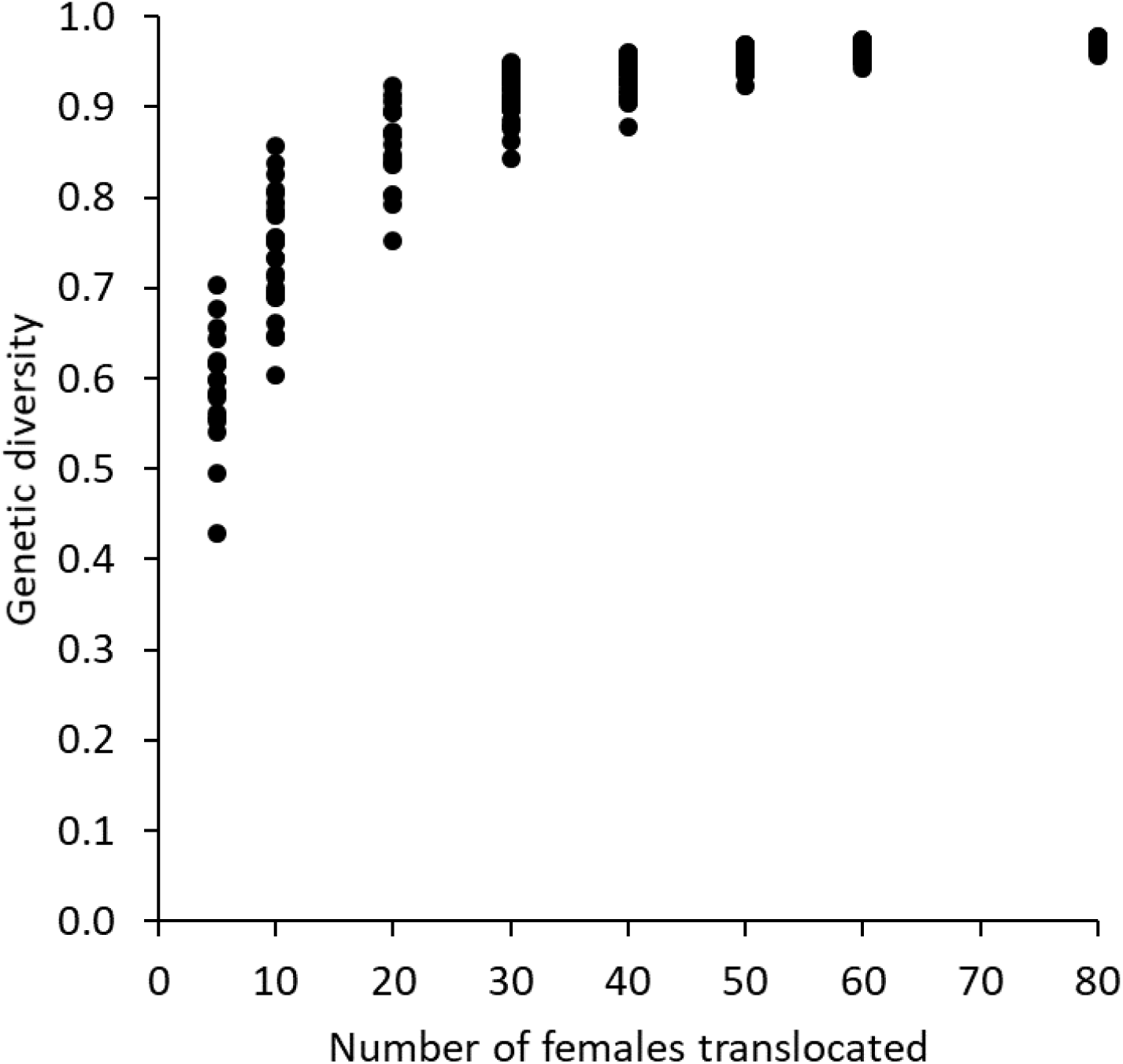
Gene diversity in final extant populations increased as the number of released females increased in 100-year population projections simulating 198 translocations of wild giraffes. Simulations used population growth rates: λ = 1.001, 1.010, 1.024; and genetic load number of lethal equivalents: LE = 2.5, 6.29, 12.6.

Sensitivity analysis revealed demographic parameters of first-year survival and adult female reproductive rate were the most influential on long-term viability metrics of PE and GD (Table 3). Changes in first-year mortality and reproductive rate both increased PE > 10% and reduced GD > 10%. Increasing the age of first reproduction by 1 year resulted in a 1% increase in PE and 2% reduction in GD, and all other parameter changes resulted in < 1% reduction in PE and ≤ 2% reduction in GD (Table 3).

**Table 3.** Sensitivity analysis of demographic and genetic load parameters for simulations of giraffe translocations where each parameter was proportionally reduced ∼25%. ΔPE is change in probability of extinction relative to the reference simulation, ΔGD is change in genetic diversity relative to the reference simulation.

| Parameter | Value | ΔPE | ΔGD |
| --- | --- | --- | --- |
| First-year mortality | 63% | 0.114 | -0.106 |
| Reproductive rate | 38% | 0.102 | -0.105 |
| Age of first reproduction | M=3, F=4 | 0.009 | -0.024 |
| Second-year mortality | 25% | 0.002 | -0.020 |
| Adult mortality | 6% | 0.000 | -0.009 |
| Genetic load lethal equivalents | 7.90 | 0.003 | -0.006 |

## DISCUSSION

Despite the potentially low chance of success (Fischer and Lindenmeyer 2000), translocations continue to be used as a major tool in endangered species management. Our analyses evaluated the likely result of a giraffe translocation under reasonable demographic and genetic assumptions based on published literature. We considered a population to be minimally demographically viable when it has a 5% or lower probability of extinction during a 100-year period, and minimally genetically viable when it has a 20% or lower decrease in genetic diversity compared to the original wild population over the same period (IUCN 2013). These minimum criteria for viability of an isolated population were achieved when 20 females and 2 males were released, assuming demographic parameters for a moderate population growth rate (λ = 1.010), and a moderate amount of inbreeding genetic load (lethal equivalents: LE = 6.29). Under more conservative demographic rates and higher genetic loads, the minimum release population for 95% viability at 100 years was 30 females and 3–6 males. This latter scenario also managed to preserve at least 80% of the genetic diversity of the source population. Population projections from our scenarios illuminate the marked impact of population growth and genetic load on the long-term viability and genetic diversity. Our study thus shows how uncertainty in these parameters can affect the minimum number of founders and sex ratios of animals required to be released to successfully establish a new population.

Few studies have thoroughly monitored source and translocated populations for changes in genetic parameters such as heterozygosity, allelic richness, and rate or level of inbreeding (Groombridge et al. 2012, Puckett et al. 2014). Ideally, genetic variability should be assessed in source populations in advance of translocations to use genetic information to guide translocation plans and to provide a contrast to post-translocation genetic studies (Rocamora and Richardson 2003, Biebach and Keller 2010). This is particularly important if there is reason to assume that the source population already has low genetic diversity, for example, because it was established through translocations or because it was a long-isolated population of only a few hundred individuals. Consideration should also be given to the issues that can arise when using individuals for reintroduction from captive stocks where genetic adaptation to captivity may have occurred (Montgomery et al. 2010, Robert 2009), which becomes more likely with more generations in captivity, but should have been partially mitigated if the population has been managed to minimize inbreeding (Marsden et al. 2013, Frankham 2010).

During the “founders’ years” from year 2 to 23 in all our projections, initial population growth will likely be positive if 2 or more females are translocated. However, this initial growth is a transient effect of the young-skewed age distribution of the founding population, and very few translocations of small (<20 females) founding populations will be viable in the long term in isolation due to a lack of genetic diversity (**Fig 1**). Increasing the number of females in the initial release had a strong positive effect on mean annual stochastic rate of population increase, and final genetic diversity. A 0.01 probability of extinction was achieved assuming moderate demographic rates and moderate genetic load if the founder population included 30 females and 3 males. In situations where the translocation destination has few or no large predators, and asymptotic growth rates could approach 1.024, then viable populations could be achieved with releases of a minimum of 20 females and 2 males that are all unrelated. These minimum founding populations will result in a 20% loss of genetic diversity, so if genetic diversity considerations are paramount and additional translocations at a later stage are unlikely, then much larger founding populations should be attempted.

Note that this is not due to a lack of genetic diversity captured by the founding population, but due to the rapid loss of genetic diversity in the first years when the population is still small. A population that is twice as large, loses genetic diversity at half the speed. This highlights the importance of ensuring that a reintroduced population can grow as fast as possible in this first stage. Population growth could be facilitated, for example, by providing supplemental water or food, or veterinary care, especially for individuals with under-represented genes. This may seem to counteract natural selection, but selection works inefficiently on small populations (Margan et al. 1998). Instead, maintaining genetic diversity will maximize the material available for selection once the population has reached a stable size of at least several hundreds of individuals.

Also, the loss of genetic diversity can be mitigated by immigration of a few individuals from another population every generation, with the exact number of migrants being dependent on the situation (Wang et al. 2004, Puckett et al. 2014). If there are natural corridors, this exchange may happen naturally. For fenced or isolated areas, this will require periodic translocations of individuals. Either way, this carries the risk of spreading diseases.

Rules of thumb for translocations such as we presented here should not be used as a substitute for site-specific monitoring and analyses, but they can be useful in early decision-making and feasibility analyses. Ideally, every translocation should be conducted under an adaptive management framework with continuous pre- and post-translocation monitoring of the source and destination populations and annual updates to PVA analyses to inform structured decision-making (McCarthy et al. 2011, Williams and Brown 2014). The use of continuous monitoring, formal and transparent decision-making, and data-driven adaptive management cycles will generate the necessary data to estimate demographic parameters, identify prospective subject animals, predict population viability, and make informed management decisions. The true demographic rates of any translocated population should be estimated annually and compared with assumptions made during planning. As our sensitivity analysis showed, if demographic rates such as juvenile survival or reproductive rates are below critical levels, the translocated population will be considerably less viable than was predicted.

The ideal female age class for translocations in terms of population growth rate is the youngest age class that can immediately begin reproducing, but adults are logistically challenging to move due to their large body size. All of our population projections used the logistically more tractable age class of 2-year-olds for translocation, as this age class is often chosen for translocations. Translocating juveniles means there is an inevitable initial post-release decline in population size as animals are subject to mortality before reaching reproductive age. Sensitivity analyses indicated ages of first reproduction above our very optimistic values of 3 for females and 2 for males can adversely affect rate of population increase and long-term viability.

Fewer males than females are necessary in translocations because the male contribution to population viability is mostly confined to their genetic contributions, and equal offspring sex ratios will greatly increase the number of adult males within 1 generation. Our projections assumed all males were in the breeding pool and had equal probability of mating. The details of giraffe mating success are largely unknown for wild populations, but there is evidence that dominant males may monopolize mating opportunities potentially resulting in a skewed mating success among existing males (Pratt and Anderson 1982; 1985). Similarly, female mate choice could result in a skewed mating success among males (Andersson 1994). Either of these situations would reduce the pool of breeding males thereby reducing the genetic diversity passed to subsequent generations, leading to a smaller effective population size, lower genetic diversity, and possibly higher inbreeding depression.

Future research to inform translocation PVA analyses should focus on two critical data gaps. First, research should aim to quantify inbreeding depression in wild or semi-wild populations that have and have not experienced a population bottleneck, and to quantify genetic parameters in translocated versus source populations over time. Second, research should determine whether assumptions of panmictic breeding and equal breeding success of all males are accurate. This is important because skewed mating success will reduce the effective population size, and worsen the impact of inbreeding, i.e. result in more severe inbreeding depression. Published accounts of translocation outcomes including annual demographic rates of age-specific survival and reproduction, genetic diversity, and long-term population viability are also needed as giraffe translocations are typically not well documented (Winter et al. 2019).

## ACKNOWLEDGEMENTS

We thank two anonymous reviewers and the associate editor who improved this manuscript.

